# Dynamic BMP10 Release Reflects Atrial Fibrillation Burden in Human Atrial Engineered Heart Tissue

**DOI:** 10.64898/2026.08.20.746063

**Authors:** L. von Hacht, T. Meier, J. Ridder, J. Schrapers, A.K. Afflerbach, M.N. Hirt, A. Hansen, P. Kirchhof, T. Eschenhagen, J. Stenzig, L. Fabritz, L.C. Sommerfeld

**Affiliations:** University Center of Cardiovascular Science (UCCS), University Medical Center Hamburg-Eppendorf (UKE), Hamburg, Germany; German Center for Cardiovascular Research (DZHK), partner site North, Hamburg, Germany; Department of Cardiology, University Heart & Vascular Center, UKE, Hamburg, Germany; Institute of Experimental Pharmacology & Toxicology, UKE, Hamburg, Germany; Bioinformatics Core, University Medical Center Hamburg-Eppendorf, Hamburg, Germany; Cardiovascular Sciences, University of Birmingham, Birmingham, UK

**Keywords:** biomarker, optogenetic pacing, disease modeling

## Abstract

**Background:** Atrial fibrillation (AF) burden is increasingly recognized as a determinant of clinical risk. Currently, AF burden can only be estimated using long-term rhythm monitoring. Bone morphogenetic protein 10 (BMP10) is a protein secreted from cardiac atria associated with AF and AF-related complications. This study evaluated whether BMP10 concentrations are associated with AF burden in a human atrial model: atrial engineered heart tissue (aEHT).

**Methods:** Human induced pluripotent stem cell-derived atrial cardiomyocytes were cast into atrial engineered heart tissues (aEHTs). To mimic AF burden, mature aEHTs were optogenetically-paced at a high rate of 4 Hz, either intermittently for 4 hours every 2 days (∼10% burden) or continuously for 24 hours per day (100% burden). After 18 days of high-rate pacing intervention, 7 days of recovery without pacing followed. BMP10 release was quantified by ELISA and contractile function was assessed by video analysis. EHT transcriptional remodeling in response to mimicked AF burden was assessed by RNA sequencing and the effect of recovery was analyzed by qPCR.

**Results:** High-rate optogenetic pacing mimicking AF lead to a dynamic, burden-dependent BMP10 release: BMP10 concentrations in the medium were increased by intermittent optogenetic pacing (∼10% burden) and highest under continuous optogenetic pacing (100% burden). BMP10 release declined toward control levels during recovery. Contractile dysfunction was most impaired after continuous pacing and showed only partial recovery within 1 days after pacing cessation. RNA sequencing revealed distinct burden-dependent transcriptional states. Pacing-regulated transcripts were related to BMP/TGFβ signaling, atrial identity, calcium handling, contractile phenotype, and electrophysiological remodeling. After recovery, *BMP10* mRNA expression remained elevated despite normalization of BMP10 protein release.

**Conclusions:** AF burden dynamically regulates BMP10 release and functional and molecular remodeling in human aEHTs. BMP10 release depicts a secreted protein-based readout of current or recent atrial high-rate stress, whereas persistent transcriptional changes indicate molecular memory of prior AF burden. These findings support BMP10 release as a burden-sensitive AF biomarker.

**Graphical Abstract:** 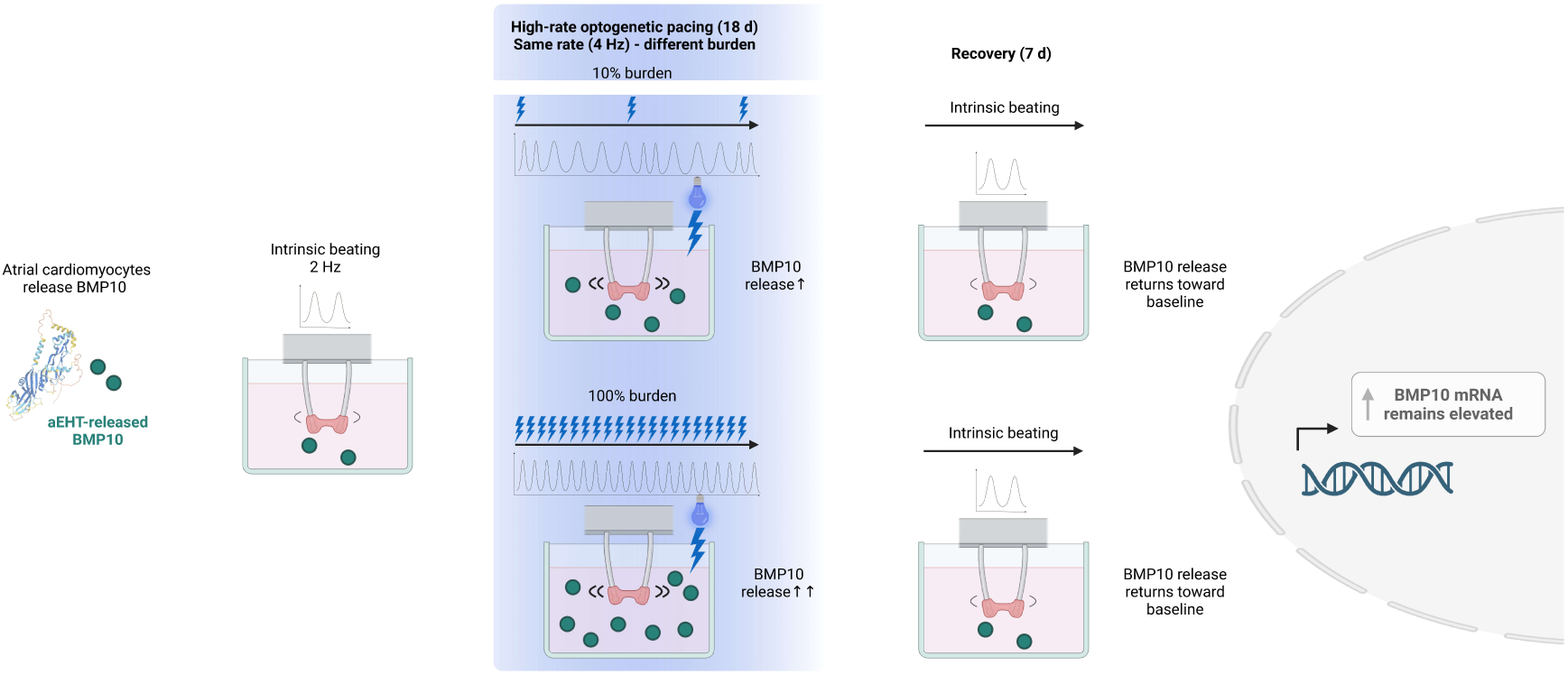

**Clinical Perspective:**

- The secreted bone morphogenetic protein 10 (BMP10) is an emerging blood biomarker associated with atrial fibrillation (AF) and its complications.
- Continuous atrial high-rate pacing leads to BMP10 release in atrial engineered heart tissues (aEHTs).
- BMP10 release and *BMP10* mRNA expression mirrors varying AF burden in aEHT and even ∼10% burden leads to significantly increased *BMP10* mRNA expression and protein release.
- After returning to normal rate, BMP10 release decreases to baseline within 7 days, while *BMP10* mRNA expression shows a delayed response.
- BMP10 may qualify as an AF burden-sensitive, circulating biomarker for AF and burden-associated outcome prediction.

## Introduction

Atrial fibrillation (AF) affects approximately 60 million individuals worldwide and is associated with increased risk of stroke, heart failure, and death. The secreted and atrial-specific growth factor bone morphogenetic protein 10 (BMP10) is associated with incident, prevalent, and recurrent AF, as well as with AF-related complications including stroke.^1,2^ Furthermore, combining BMP10 with other circulating biomarkers improves AF detection.^3^ Elevated BMP10 concentrations are associated with left atrial endomysial fibrosis in patients undergoing cardiac surgery.^4^ These results suggest that BMP10 expression and release could be influenced by atrial rate and that BMP10 might reflect structural atrial remodeling, or contribute to it, in addition to acute rate-dependent release.

Recent clinical trials^5^, including EAST-AFNET 4, ARTESIA and NOAH-AFNET 6 in device-detected AF^6,7^ and controlled trials of anticoagulation after AF ablation^8,9^, suggest that stroke risk is modulated by AF burden, reflecting the time spent in AF. Stroke risk without anticoagulation is lower in patients with device-detected AF and a lower AF burden than in patients with persistent and permanent AF, with paroxysmal AF showing intermediate rates. Subsequently, a clinical dose-response relationship between AF burden and stroke risk has been established.^10^ Quantifiable estimates of AF burden could therefore improve anticoagulation decisions and AF screening strategies. Repeated BMP10 measurements track current AF rhythm status, and elevated BMP10 levels over time further refine the associations with ischemic stroke, heart failure hospitalization, and all-cause mortality^11^, suggesting that BMP10 dynamics may reflect changing AF burden. In addition, a recent exploratory analysis of the EAST-AFNET 4 Biomolecule Study found broadly consistent benefit of early rhythm control across most circulating biomolecules, while categorical analyses identified a nominal interaction signal for BMP10, suggesting a potentially attenuated treatment effect in patients with very low BMP10 concentrations.^12^

Complementing these clinical observations, continuous high-rate optogenetic pacing increases concentrations of released BMP10 from atrial engineered heart tissue (aEHT).^13^ Encouraged by these findings, we now used the human aEHT model to mechanistically investigate how AF burden regulates BMP10 release and aEHT function. We exposed human aEHTs to defined intermittent and continuous optogenetic high-rate pacing regimens, mimicking different AF burdens and assessed BMP10 release, contractile function and transcriptional remodeling, as well as recovery from atrial high rate episodes.

## Methods

### Generation, Culture, and Contractile Analysis of Human Atrial Engineered Heart Tissues

Human atrial engineered heart tissues (aEHTs) were generated and analyzed following previously established protocols with minor modifications.^14–16^ Human induced pluripotent stem cells from the ERC001 (https://hpscreg.eu/cell-line/ERCi001-A) line, an in-house quality-controlled cell line established at the University Medical Center Hamburg-Eppendorf, were differentiated into atrial cardiomyocytes using established atrial differentiation protocols.^14,16^ For aEHT generation, 1 × 10⁶ human iPSC-derived atrial cardiomyocytes were embedded in a fibrin-based hydrogel and cast into 24-well silicone post-based EHT molds.

After casting, aEHTs were maintained under previously-published standard conditions with medium exchange three times per week and allowed to mature for 39 days before initiation of optogenetic high-rate pacing.^13,17^ Contractile function was assessed longitudinally by automated video-optical analysis of post deflection using the CTMV software (EHT Technologies). Spontaneous beating frequency and contractile force were determined in the absence of external stimulation during maturation, pacing, and recovery period.

### Optogenetic High-Rate Pacing

To enable optogenetic stimulation, aEHTs were transduced during casting with adeno-associated virus serotype 6 encoding the light-gated nonselective cation channel CheRiff2.0 under control of a cardiomyocyte-specific cTNT promoter. Blue light stimulation was delivered using a custom-built LED-based pacing platform. Unless stated otherwise, optical stimulation was performed at 470 nm with 40 ms light pulses (0.12 mW/mm²).^13,17^

On day 39 after casting, aEHTs were assigned to one of three experimental groups: unpaced control, intermittent atrial high-rate optogenetic pacing, or continuous atrial high-rate optogenetic pacing - mimicking ∼10% and 100% AF burden, respectively. Control tissues were maintained at their intrinsic spontaneous beating frequency of approximately 120 beats per minute. Intermittently paced aEHTs were stimulated at 4 Hz for 4 hours followed by 44 hours without optogenetic pacing, corresponding to a burden of approximately 10% over each 48-hour cycle. Continuously-paced aEHTs were stimulated at 4 Hz for 24 hours per day, corresponding to a 100% burden. The 10% burden threshold was chosen based on clinical observations. This threshold has been shown to occur commonly in patients with device-detected AF^7^ and has also been shown to negatively impact outcome in patients.^10^ Optogenetic pacing was performed for 18 days, followed by a 7-day recovery period during which all aEHTs were maintained without external stimulation and allowed to beat spontaneously.

### Collection of Conditioned Medium and BMP10 Quantification

Conditioned medium was collected at each medium exchange during maturation, pacing, and recovery. At the time of media change, 1 mL of conditioned medium was collected from each well, snap-frozen in liquid nitrogen, and stored at −80°C until analysis. BMP10 concentrations in conditioned medium were quantified as described previously^13^ using the Human BMP-10 ELISA Kit PicoKine EK1808 (BosterBio, Pleasanton, CA, USA) according to the manufacturer’s instructions. Samples were diluted 1:10-1:50 in PBS. Absorbance was measured using a Multiskan SkyHigh Microplate Spectrophotometer (Thermo Scientific).

### RNA Isolation and Quantitative PCR

Total RNA was isolated from snap-frozen aEHTs using TRIzol reagent followed by chloroform phase separation and purification with the RNeasy Mini Kit (QIAGEN), including on-column DNase digestion to reduce genomic DNA contamination. RNA was eluted in 15 µL RNase-free water, and concentration and purity were assessed using a Multiskan SkyHigh Microplate Spectrophotometer.

Complementary DNA was synthesized from 200 ng total RNA using the High Capacity cDNA Reverse Transcription Kit (Thermo Fisher Scientific). Quantitative PCR was performed using PowerTrack SYBR Green Master Mix (Thermo Fisher Scientific) on a QuantStudio 5 Real-Time PCR Instrument (Thermo Fisher Scientific). Relative gene expression was calculated using the ΔCt method and normalized to the geometric mean of *POLR2A* and *PUM1*. Primer sequences are provided in Supplementary Table 1.

### Bulk RNA Sequencing and Bioinformatic Analysis

For transcriptomic profiling, total RNA from control, ∼10% burden, and 100% burden aEHTs harvested directly after the optogenetic pacing period was subjected to bulk RNA sequencing. Raw reads were processed with fastp (v0.23.2) to remove adapter-derived and low-quality sequences, including correction of mismatched base pairs in overlapping regions using the -- correction option. Reads were aligned to the human reference assembly GRCh38.110 using STAR (v2.7.10a). Gene-level counts were generated using the STAR GeneCounts option (v2.7.10a). Batch effects were corrected using ComBat-Seq implemented in the sva package (v3.54.0). Differential expression analysis was performed with DESeq2 (v1.42.0) using batch-corrected counts. Genes were considered significantly differentially expressed if the absolute log2-transformed fold change was ≥1 and the false discovery rate was ≤0.05.

Upon publication, data will be made publicly available at https://www.ncbi.nlm.nih.gov/geo/.

### Statistical Analysis

All biochemical assays were run in a blinded fashion. Data are presented as mean ± SEM unless stated otherwise. Comparisons between more than two groups were performed using one-way or two-way ANOVA, corrected for multiple comparisons (Tukey’s post-hoc test was applied for all pairwise comparisons). Longitudinal data were analyzed using repeated-measures ANOVA. Adjusted *p* values are reported for multiple comparisons. Statistical analyses and visualization were performed using GraphPad Prism version 10 and R version 4.5.2. A two-sided *p* value or adjusted *p* value <0.05 was considered statistically significant. Volcano plots were generated in R (version 4.5.2) using ggplot2 and ggrepel by plotting log2 fold change against −log10-transformed nominal *p* value. Genes were colored using thresholds of FDR ≤ 0.05 and |log2FC| ≥ 1 and selected significant genes together with *BMP10* were annotated.

## Results

### Atrial High Rates Induce Burden-Dependent Contractile Dysfunction and Increased BMP10 Release

After maturation, aEHTs displayed stable spontaneous beating at approximately 120 beats per minute and generated contractile force of approximately 0.33 mN. ELISA analysis of conditioned medium also showed a plateaued, reliable and consistent BMP10 release across all aEHTs. Transduced aEHTs reliably followed optical stimulation at 4 Hz (240 beats per minute), while maintaining contractile force during acute pacing assessment (Supplementary Figure 1). To model different AF burdens, mature aEHTs were subjected to intermittent 4 h/2 d optogenetic high-rate pacing, corresponding to a ∼10% burden, or continuous 24 h/day optogenetic high-rate pacing, corresponding to a 100% burden. 100% burden caused an immediate and sustained reduction in contractile force compared with unpaced controls and ∼10% burden aEHTs (Figure 1A). In contrast, force of aEHT subjected to only ∼10% burden remained closer to unpaced controls throughout the optogenetic pacing period. After cessation of optogenetic pacing, contractile force partially recovered in the 100% burden aEHTs within 7 days, indicating at least partial reversibility of optogenetic pacing-induced contractile dysfunction.

**Figure 1.**
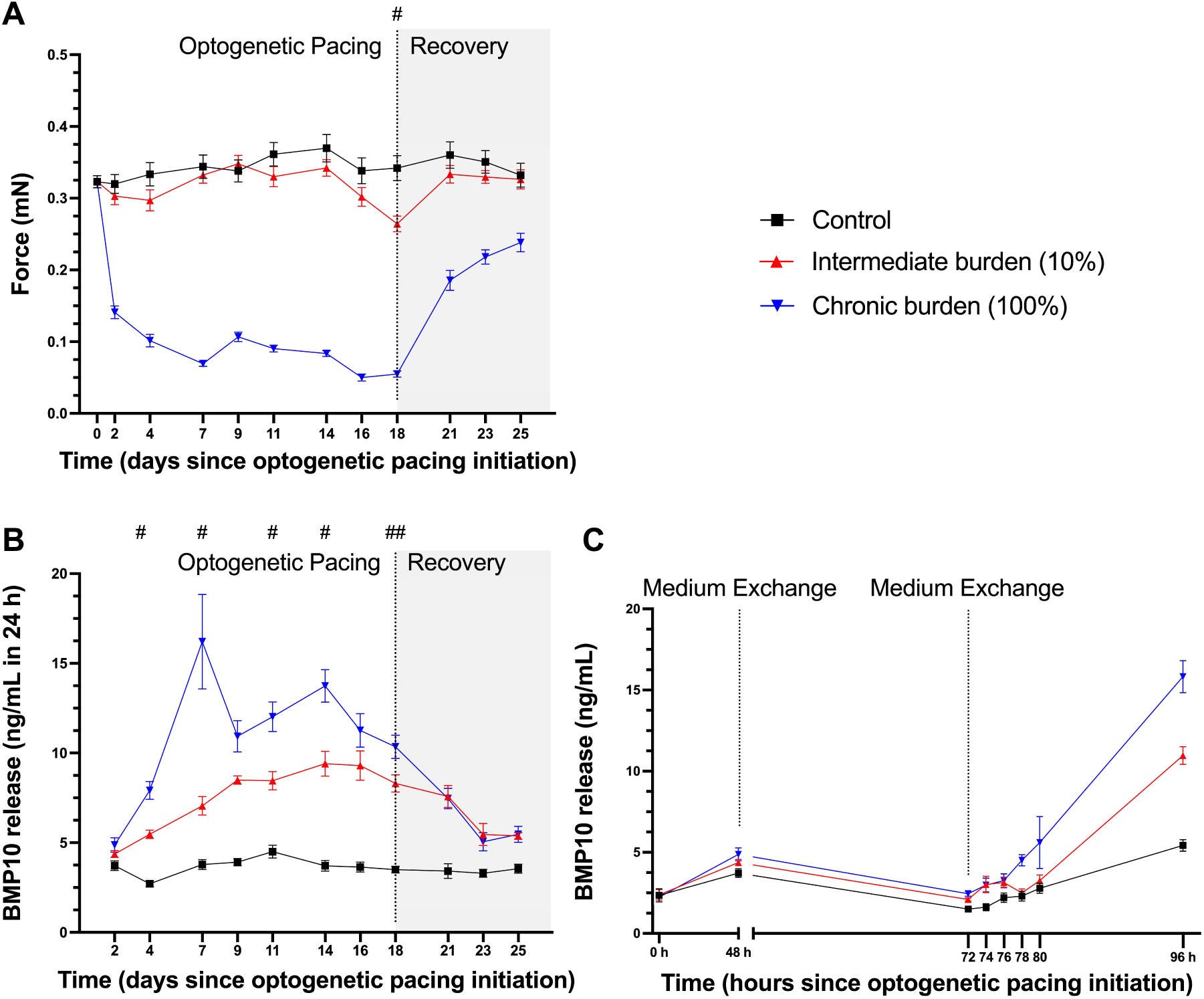
Optogenetic high-rate pacing at different burden induces burden-related contractile dysfunction and BMP10 release. **(A)** Contractile properties, assessed by video optical recordings in the absence of stimulation, of control, ∼10% and 100% burden aEHTs during optogenetic pacing and recovery (n=16-22/group). **(B)** BMP10 concentration in medium during atrial high-rate optogenetic pacing and recovery (n=7-8/group). **(C)** BMP10 concentration in medium from the initiation (0 h) of high-rate optogenetic pacing over 96 hours. Complete medium exchange took place at 48 h and 72 h (n=7-8/group). The 96 h time point is equivalent to 4 days since pacing initiation. Two-way ANOVA followed by Tukey’s multiple comparisons test (# indicating all pairwise comparisons are significant, ## indicating 100% vs control and ∼10% vs control are significantly different).

**Figure 2.**
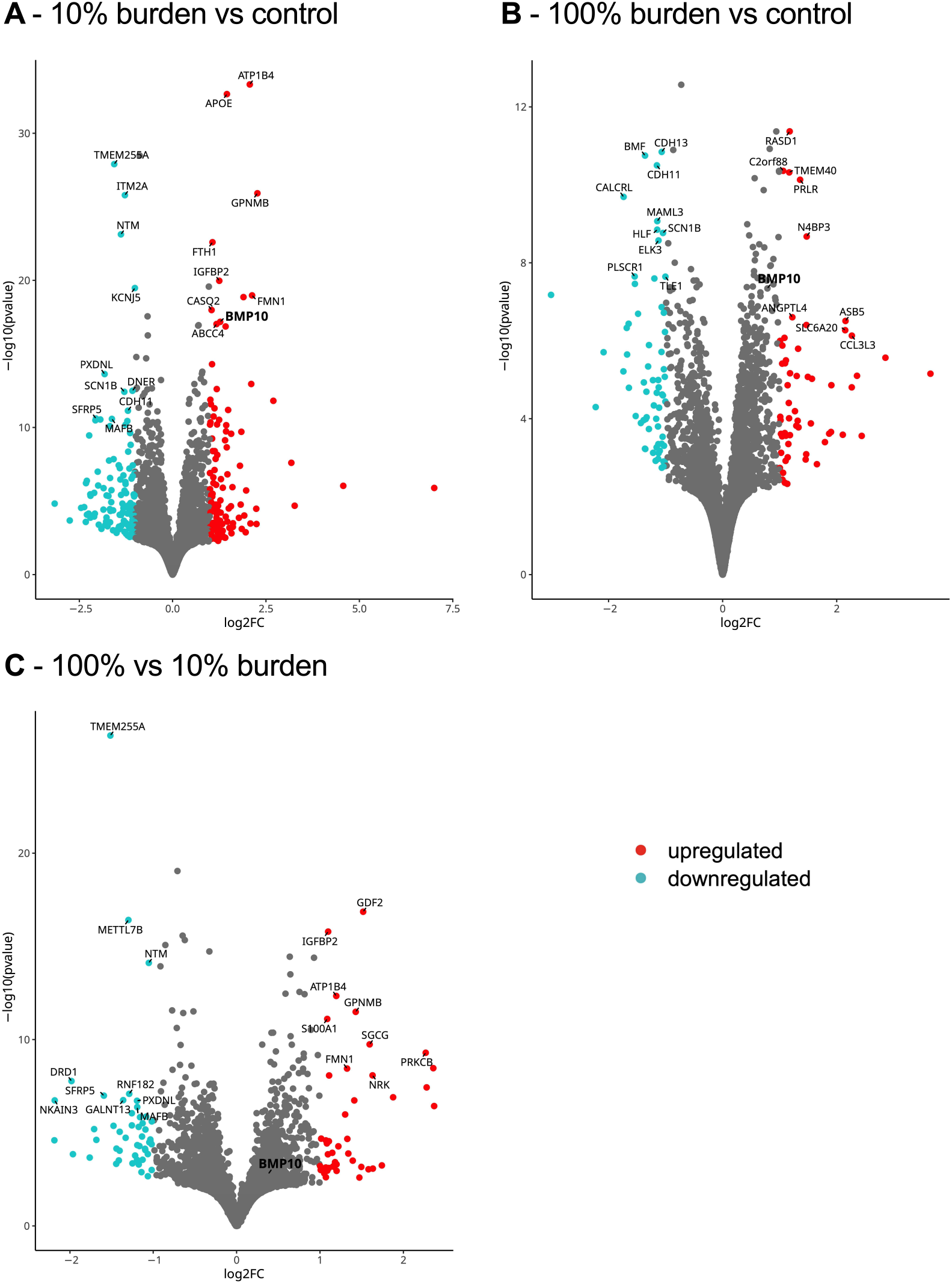
*BMP10* as a prominent transcript affected by high-rate pacing. Volcano plots of pairwise comparisons of bulk RNA sequencing differential expression analysis of aEHTs subjected to different burden for 18 days. **(A)** ∼10% burden versus control. **(B)** 100% burden versus control. **(C)** 100% burden versus ∼10% burden. Red points indicate upregulated genes, and cyan points indicate downregulated genes according to the predefined significance and log2FC thresholds. N=5/group

BMP10 release increased in response to optogenetic high-rate pacing in a burden-dependent manner. Compared with unpaced controls, both ∼10% and 100% burden increased BMP10 concentrations in conditioned medium, with the highest concentrations observed in the aEHTs subjected to 100% burden (Figure 1B). BMP10 release increased during the first week of optogenetic pacing, remained elevated during ongoing optogenetic pacing, and declined after cessation of the pacing. During recovery, BMP10 concentrations approached control levels within 5-7 days, indicating a dynamic and reversible secretory response to burden.

To further characterize acute BMP10 release kinetics, conditioned medium was collected at defined intervals after the last complete medium exchange on day 3 of the ongoing high-rate optogenetic pacing protocol. Released BMP10 accumulated progressively over 48 hours, with highest accumulation under 100% burden. In the ∼10% burden group, BMP10 continued to accumulate despite the intermittent pacing pattern and periods without high-rate pacing stimulation, indicating sustained release beyond the active atrial high-rate window of 4 hours (Figure 1C). In both burden groups, the BMP10 release continued to increase after 96 hours (day 4) and reached the highest concentrations in the 100% burden group after 7 days (Figure 1B).

### *BMP10* is a Prominent Burden-Associated Transcript

Analysis of the global transcriptional response to varying degrees of burden over 18 days revealed consistent *BMP10* upregulation. *BMP10* was among the prominent upregulated transcripts after atrial high-rate pacing with both, intermediate (∼10%) and high (100%) burden. *BMP10* was among the top 10 most upregulated (statistical significance and above log2FC threshold) genes with 100% burden, and already showed a significant, but less pronounced upregulation with ∼10% burden compared to unpaced controls.

### Atrial High-Rate Burden Regulates BMP/TGFβ, Contractile, Calcium-handling, and Channel Gene-Related Transcripts

Pacing-induced expression changes varied with degree of burden (Figure 3). *BMP10* expression increased with higher burden, with highest expression observed after 100% burden for 18 days. Genes associated with atrial identity and BMP/TGFβ-related signaling showed no obvious burden-dependent regulation. Compared to control, *PITX2*, a prominent atrial transcription factor, and *ENG/Endoglin*, a co-receptor for BMP signaling, were expressed at lower concentrations with 100% burden pacing only, while *ID4*, part of the downstream BMP-cascade, were significantly less expressed with ∼10% and 100% burden. However, none of these transcripts showed significant expression differences between ∼10% and 100% burden. Changes in calcium-handling and contractile remodeling were reflected by higher *CASQ2* expression after 100 % burden optogenetic pacing, and a higher *MYH7*/*MYH6* expression ratio, indicating an atrial high rate-associated shift in contractile gene expression. Downregulation of selected ion channel-related transcripts compared to control indicated selective remodeling, with burden-mirroring lower expression of *CACNA1G*, *KCNJ5*, and *SCN1B,* while *SCN5A* showed no expression difference between burden groups. Additionally, transcripts showing non-linear expression response to varying burden were also observed (Supplementary Figures 2), suggesting a complex relationship rather than a unidirectional, dose-response effect. Varying burden levels also produced distinct enrichment patterns across a broad range of gene sets (Supplementary Figure 3).

**Figure 3.**
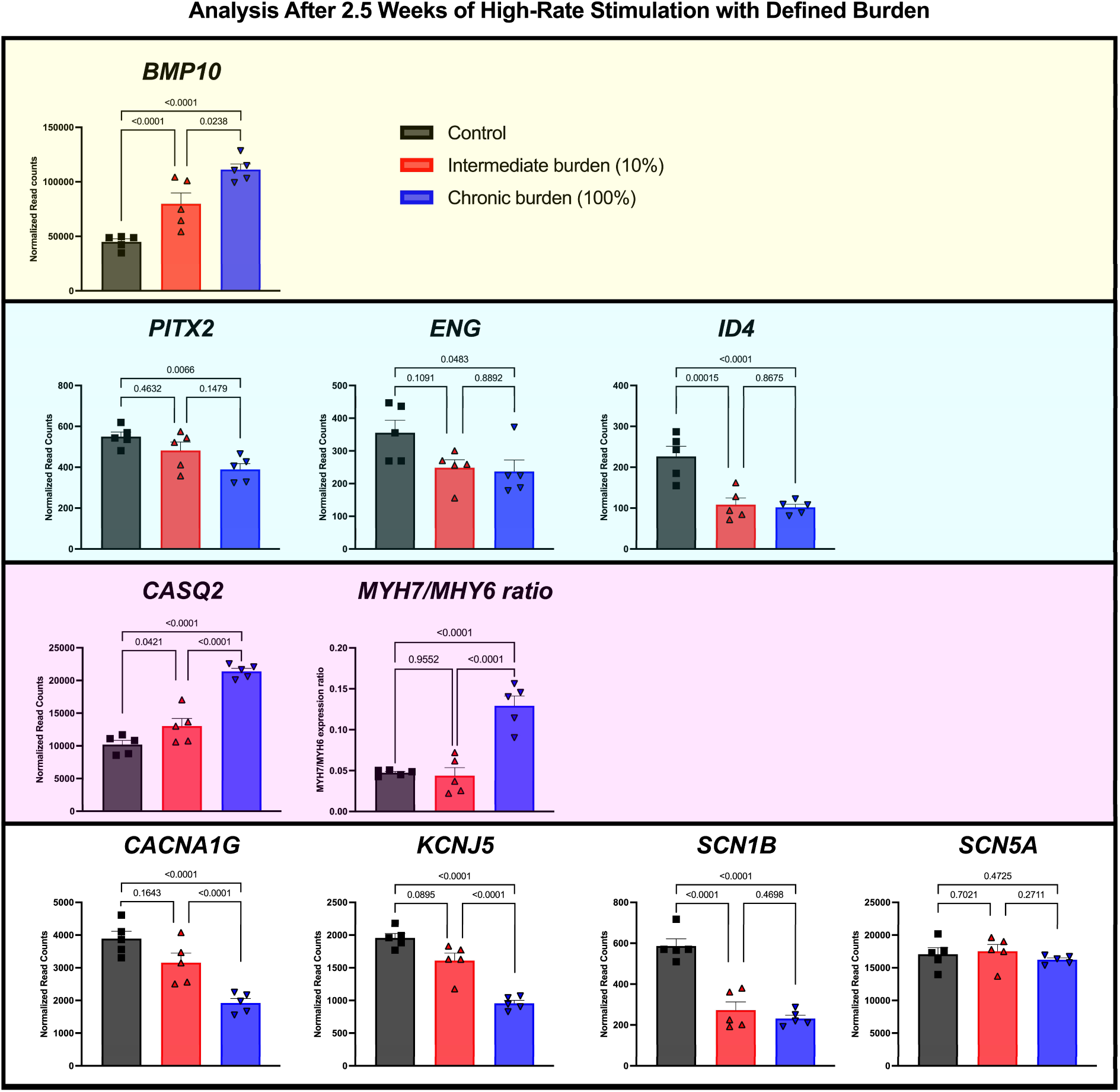
Burden-dependent *BMP10* upregulation and further transcriptional remodeling in atrial high-rate optogenetically-paced aEHTs. Bulk RNA sequencing was performed on aEHTs harvested directly after the 18-days optogenetic pacing period. *BMP10* (faint yellow background), selected *BMP10*-related transcripts (cyan background), calcium-handling and contractile remodeling (magenta background) as well as ion channel transcripts (clear background) are shown across control, ∼10% and 100% burden groups (n=5/group). Adjusted *p* values are shown. The *MYH7*/*MYH6* ratio was calculated from normalized expression values per sample and analyzed separately using one-way ANOVA with adjusted *p* values. Gene symbols: *BMP10*, bone morphogenetic protein 10; *PITX2*, paired-like homeodomain transcription factor 2; *ENG*, endoglin; *ID4*, inhibitor of DNA binding 4; *CASQ2*, calsequestrin 2; *MYH7*, myosin heavy chain 7; *MYH6*, myosin heavy chain 6; *CACNA1G*, calcium voltage-gated channel subunit alpha1 G; *KCNJ5*, potassium inwardly rectifying channel subfamily J member 5; *SCN1B*, sodium voltage-gated channel beta subunit 1; *SCN5A*, sodium voltage-gated channel alpha subunit 5.

### High-Rate-Induced Increase in *BMP10* Expression Persists 7 Days after Cessation of High-Rate Pacing

To assess whether atrial high-rate-associated transcriptional changes persisted after cessation of high-rate optogenetic pacing, targeted qPCR was performed after 7 days of recovery in control, ∼10% and 100% burden aEHTs. *BMP10* mRNA expression remained elevated after recovery in both burden groups compared with control aEHTs (Figure 4A). Among selected remodeling-associated transcripts, *PITX2*, *ENG*, and *SCN1B* no longer showed significant differences between groups after recovery. *CASQ2* remained higher in burden-subjected aEHTs and showed the highest expression in response to previous 100% burden. *MYH7* expression was higher after 100% burden compared with ∼10% burden, whereas *MYH6* expression was not significantly different between the groups. Accordingly, the *MYH7*/*MYH6* ratio was higher after recovery from 100% burden pacing. *CACNA1G* expression was lower after recovery from ∼10% burden compared with unpaced controls, while *KCNJ5* expression was lower after recovery from 100% burden compared with unpaced controls.

**Figure 4.**
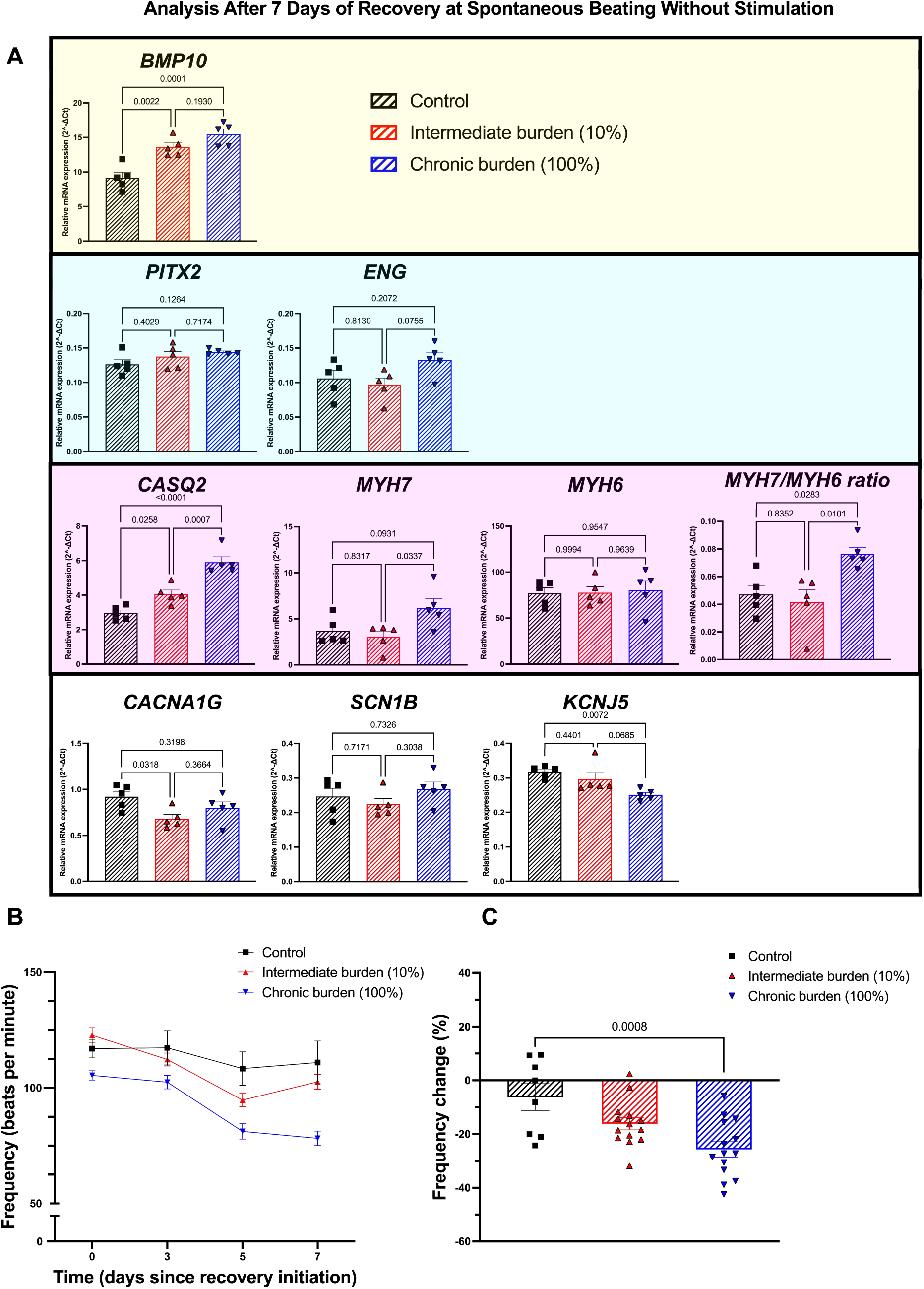
***BMP10* upregulation persists amid reduced spontaneous beating frequency with selective transcriptional normalization after 7 days of recovery** (A) Targeted qPCR analysis of selected remodeling transcripts in human atrial EHTs after 18 days of high-rate pacing (unpaced control, ∼10% or 100% burden) and additional 7 days of recovery without optogenetic pacing in any group. Relative mRNA expression of *BMP10* (faint yellow background) and transcripts related to BMP10 signaling (cyan background), calcium handling and contractile remodeling (magenta background) as well as ion channels (clear background) are shown (n=5/group). Ordinary one-way ANOVA followed by Tukey’s multiple comparisons test. Adjusted *p* values are shown. **(B)** Contractile properties were assessed by video optical recordings. Intrinsic beating frequency of human aEHTs during 7 days of recovery after cessation of high-rate optogenetic pacing (n=8-14/group). **(C)** Change of intrinsic beating frequency from the day of recovery initiation to day 7 of recovery. Ordinary one-way ANOVA followed by Tukey’s multiple comparisons test. Adjusted *p* values are shown (n =8-14/group). Gene symbols: *BMP10*, bone morphogenetic protein 10; *PITX2*, paired-like homeodomain transcription factor 2; *ENG*, endoglin; *CASQ2*, calsequestrin 2; *MYH7*, myosin heavy chain 7; *MYH6*, myosin heavy chain 6; *CACNA1G*, calcium voltage-gated channel subunit alpha1 G; *SCN1B*, sodium voltage-gated channel beta subunit 1; *KCNJ5*, potassium inwardly rectifying channel subfamily J member 5.

In addition to incomplete transcriptional recovery and impaired contractile recovery, aEHTs subjected to atrial high rates displayed a burden-dependent slowing of intrinsic beating frequency during recovery, with the strongest depression in beating frequency observed in aEHT revering from 100% burden (Figure 4B and 4C).

## Discussion

### Main Findings

●**BMP10** expression and release reflect AF burden, mimicked by optogenetic atrial high-rate pacing, in mature aEHTs.
●**Reversibility**: BMP10 release declines after high-rate pacing discontinuation, marking BMP10 as a dynamic, reversible protein-level indicator of current/recent atrial stress.
●**Molecular memory:** During recovery from atrial high-rate episodes, *BMP10* transcriptional expression stays elevated despite normalized BMP10 release. Regulatory/remodeling transcripts show divergent recovery, and intrinsic spontaneous beating frequency declines in a burden-dependent manner, suggesting persistent transcriptional changes may represent a lasting molecular memory beyond the protein-level signal.

### BMP10 Release is Increased by High-Rate Pacing in an AF Burden-Dependent Manner

After maturation of the aEHTs, optogenetic pacing induced a burden-dependent phenotype characterized by reduced contractile force and increased BMP10 release. Importantly, both burden groups were stimulated at the same rate of 4 Hz, while cumulative atrial high-rate exposure differed between ∼10% and 100% burden. This design allowed cumulative atrial high-rate *exposure* (similar to “time spent in AF”), rather than frequency, to be isolated as the key experimental variable. Thus, the observation that even ∼10% burden significantly increased BMP10 release and upregulated *BMP10* transcription indicate that BMP10 regulation and release do not require 100% burden but respond already to limited recurrent high-rate exposure. While high-rate pacing at 100% burden has previously been shown to increase BMP10 release^13^, the present data demonstrate that ∼10% burden is sufficient to activate the BMP10 release response. Therefore, BMP10 concentrations emerge as a burden-sensitive, rather than merely rate-responsive atrial stress signal.

### AF Burden Induces Coordinated Transcriptional Remodeling with Distinct Recovery Kinetics

Bulk RNA sequencing placed *BMP10* within a broader atrial high-rate-associated transcriptional response. *BMP10* appeared among the most prominent upregulated transcripts after atrial high-rate exposure, including after ∼10% burden, supporting the concept that BMP10 is not only an a priori biomarker candidate but also part of the global transcriptional response to recurrent atrial high-rate exposure.

*ENG* and *ID4*, as part of the BMP-receptor and downstream signaling complex, were also regulated by AF burden, suggesting broader involvement of developmental and BMP/TGFβ-associated transcriptional programs. *ENG* has been linked to pulmonary hypertension and heart failure, but a direct link to AF has not been established.^18,19^ *PITX2* downregulation under 100% burden is relevant because *PITX2* is a central AF-susceptibility gene and has been linked to atrial gene-network regulation.^1,20^ Reduced left-atrial *PITX2* has additionally been shown to predict AF recurrence after ablation.^1^ In parallel, AF burden affected calcium-handling and contractile-remodeling transcripts, including *CASQ2* and the *MYH7*/*MYH6* ratio, consistent with stress-associated remodeling of calcium storage and contractile phenotype.^21^ Ion-channel-associated transcripts were selectively regulated, with lower *CACNA1G*, *KCNJ5*, and *SCN1B* expression, whereas *SCN5A* was not significantly altered. This pattern supports selective electrophysiological remodeling rather than broad nonspecific ion-channel downregulation and is consistent with previous links between *CACNA1G*, *KCNJ5/GIRK4*, sodium-channel β-subunits and AF-related electrophysiological remodeling.^22–24^

Recovery analyses revealed that the protein and transcript layers of the BMP10 response follow different temporal dynamics. BMP10 release declined toward control levels within 7 days after cessation of high-rate optogenetic pacing, supporting BMP10 release as a dynamic protein-level readout of current or recent atrial high-rate exposure. In contrast, *BMP10* transcript levels remained significantly elevated after 7 days of recovery, albeit with no significant differences between ∼10% and 100%, suggesting that the atrial tissue does not immediately return to its pre-stress transcriptional state. Other remodeling-associated transcripts showed divergent recovery patterns: *PITX2* was no longer significantly altered, whereas *CASQ2*, the *MYH7*/*MYH6* ratio, and *KCNJ5* remained altered in selected comparisons. Especially the persistently higher *MYH7*/*MYH6* ratio and higher *CASQ2* expression argue for a longer lasting remodeling of the contractile phenotype, similar to the situation in human atria of atrial fibrillation patients.^21,25^ In addition, intrinsic beating frequency declined in a burden-dependent manner during recovery, with the strongest effect after 100% AF burden. Together, these findings indicate that recovery is not a uniform return to baseline, but a layered process in which BMP10 release normalizes earlier while selected transcriptional and functional changes persist.

### Translational Relevance of BMP10 as a Dynamic AF Burden Biomarker Candidate

The burden sensitivity and reversibility of BMP10 release are relevant in the context of AF, where clinical risk is increasingly understood to depend not only on the presence of AF, but also on the amount and temporal pattern of atrial high-rate activity (i.e. AF burden), including stroke risk and other AF-related complications.^10,26^ Expert consensus suggests that AF burden could be a surrogate outcome in clinical trials^27^ in addition to its role in refining risk of AF-related complications.^10,26^ An atrial biomarker that responds dynamically to varying degrees of AF burden could therefore provide complementary information to binary rhythm classification. Previous and our new findings support several properties for BMP10 that would be desirable for such a biomarker candidate: atrial specificity, measurable release into conditioned medium, sensitivity to AF burden, and reversibility after cessation of atrial high-rate activity. More research is needed to characterize the dynamic release patterns of BMP10 and its function in healthy and diseased human atria.

### Strengths and Limitations

A major strength of this study is the use of a controlled human atrial EHT model to investigate BMP10 regulation under defined atrial high-rate burden. In contrast to clinical studies, where AF burden, atrial substrate, comorbidities, medication, renal clearance, and systemic factors are difficult to separate, our experimental design allowed controlled manipulation of atrial high-rate exposure (burden).

The pacing protocol employed here replicates high atrial rates but not irregular activation, conduction heterogeneity, stretch, autonomic input, inflammation, systemic neurohumoral signaling or multicellular remodeling. But although our approach is based on a simplified model, itreflects what has been observed in clinical samples, where BMP10 plasma concentrations were higher in patients with current AF than in AF patients in sinus rhythm.^13^ Although regular 4 Hz optogenetic pacing does not fully reproduce irregular clinical AF, previous work in human iPSC-derived atrial engineered myocardium showed that irregular arrhythmogenic pacing did not further increase tachypacing-induced electrical remodeling compared with regular tachypacing.^28^ Furthermore, the applied aEHT model is useful to study cardiomyocyte-intrinsic BMP10 regulation, but it does not capture the full cellular complexity of the atria.

Of note, the observed release dynamics are experimental and cannot directly define plasma concentrations, clinical thresholds, half-life, clearance, or diagnostic specificity in patients. Previous work in optogenetically tachypaced atrial-like EHTs reported increased cardiac troponin I release into the culture medium^29^, suggesting loss of cardiomyocytes in these kind of models and we cannot definitely exclude the possibility of passive release due to e.g., cell death. However, the decline of BMP10 release after optogenetic pacing cessation and the partial recovery of contractile force argue against pacing-induced cardiomyocyte loss as the sole explanation of our observations.

This study serves a first step towards a potential role of BMP10 as a marker reflecting AF-induced atrial functional and gene regulation changes in patients in a burden-dependent manner. In addition to studies in animal models and in human atria, the effects of secretion, internalization, and modification of BMP10 by the body and its organs need to be integrated into its clinical interpretation. Future studies should integrate repeated BMP10 measurements with electrophysiological phenotyping, calcium handling, multi cell-type atrial models, and clinical cohorts with quantified AF burden to determine the translational value of BMP10 for AF burden assessment and monitoring of therapeutic response.

## Conclusion

AF burden dynamically regulates BMP10 release and induces functional and molecular remodeling in human atrial engineered heart tissues. Our data provide mechanistic support for BMP10 as an AF burden-sensitive atrial biomarker candidate and support BMP10 as a protein-level readout of current or recent atrial high-rate stress. Additionally, we established human atrial EHTs as a feasible platform to study molecular mechanisms related to AF burden.

## Acknowledgement

We thank Hartwig Wieboldt, Thomas Schulze, Birgit Klampe, Grit Höppner, and Dr Vaishnavi Ameya Murukutla for expert technical support.

## References

1. Reyat JS, Chua W, Cardoso VR, Witten A, Kastner PM, Kabir SN, Sinner MF, Wesselink R, Holmes AP, Pavlovic D, et al. Reduced left atrial cardiomyocyte PITX2 and elevated circulating BMP10 predict atrial fibrillation after ablation. JCI Insight. 2020;5. doi: 10.1172/jci.insight.139179

2. Hijazi Z, Benz AP, Lindback J, Alexander JH, Connolly SJ, Eikelboom JW, Granger CB, Kastner P, Lopes RD, Ziegler A, et al. Bone morphogenetic protein 10: a novel risk marker of ischaemic stroke in patients with atrial fibrillation. Eur Heart J. 2023;44:208–218. doi: 10.1093/eurheartj/ehac632

3. Chua W, Cardoso VR, Guasch E, Sinner MF, Al-Taie C, Brady P, Casadei B, Crijns H, Dudink E, Hatem SN, et al. An angiopoietin 2, FGF23, and BMP10 biomarker signature differentiates atrial fibrillation from other concomitant cardiovascular conditions. Sci Rep. 2023;13:16743. doi: 10.1038/s41598-023-42331-7

4. Winters J, Kawczynski MJ, Gilbers MD, Isaacs A, Zeemering S, Bidar E, Maesen B, Rienstra M, van Gelder I, Verheule S, et al. Circulating BMP10 Levels Associate With Late Postoperative Atrial Fibrillation and Left Atrial Endomysial Fibrosis. JACC Clin Electrophysiol. 2024;10:1326–1340. doi: 10.1016/j.jacep.2024.03.003

5. Kirchhof P, Camm AJ, Goette A, Brandes A, Eckardt L, Elvan A, Fetsch T, van Gelder IC, Haase D, Haegeli LM, et al. Early Rhythm-Control Therapy in Patients with Atrial Fibrillation. N Engl J Med. 2020;383:1305–1316. doi: 10.1056/NEJMoa2019422

6. Healey JS, Lopes RD, Granger CB, Alings M, Rivard L, McIntyre WF, Atar D, Birnie DH, Boriani G, Camm AJ, et al. Apixaban for Stroke Prevention in Subclinical Atrial Fibrillation. N Engl J Med. 2024;390:107–117. doi: 10.1056/NEJMoa2310234

7. Kirchhof P, Toennis T, Goette A, Camm AJ, Diener HC, Becher N, Bertaglia E, Blomstrom Lundqvist C, Borlich M, Brandes A, et al. Anticoagulation with Edoxaban in Patients with Atrial High-Rate Episodes. N Engl J Med. 2023;389:1167–1179. doi: 10.1056/NEJMoa2303062

8. Kim D, Shim J, Choi EK, Oh IY, Kim J, Lee YS, Park J, Ko JS, Park KM, Sung JH, et al. Long-Term Anticoagulation Discontinuation After Catheter Ablation for Atrial Fibrillation: The ALONE-AF Randomized Clinical Trial. JAMA. 2025;334:1246–1254. doi: 10.1001/jama.2025.14679

9. Verma A, Ha ACT, Kirchhof P, Hindricks G, Healey JS, Hill MD, Sharma M, Wyse DG, Champagne J, Essebag V, et al. The Optimal Anti-Coagulation for Enhanced-Risk Patients Post-Catheter Ablation for Atrial Fibrillation (OCEAN) trial. Am Heart J. 2018;197:124–132. doi: 10.1016/j.ahj.2017.12.007

10. Becher N, Metzner A, Toennis T, Kirchhof P, Schnabel RB. Atrial fibrillation burden: a new outcome predictor and therapeutic target. Eur Heart J. 2024;45:2824–2838. doi: 10.1093/eurheartj/ehae373

11. Gkarmiris KI, Lindback J, Alexander JH, Granger CB, Kastner P, Lopes RD, Ziegler A, Oldgren J, Siegbahn A, Wallentin L, et al. Repeated Measurement of the Novel Atrial Biomarker BMP10 (Bone Morphogenetic Protein 10) Refines Risk Stratification in Anticoagulated Patients With Atrial Fibrillation: Insights From the ARISTOTLE Trial. J Am Heart Assoc. 2024;13:e033720. doi: 10.1161/JAHA.123.033720

12. Al-Taie C, Obergassel J, Borof K, Ridder J, Rillig A, Metzner A, Goette A, Magnussen C, Sinner MF, Sommerfeld LC, et al. Interactions of blood biomolecules with early rhythm control in atrial fibrillation patients: Exploratory analysis of the EAST-AFNET 4 Biomolecule Study. Europace. 2026. doi: 10.1093/europace/euag149

13. Sommerfeld LC, Schrapers J, Muller KF, Bravo-Merodio L, Siebels B, Vermeer-Stoter AMS, Pan B, Hoppner G, O’Shea C, Ridder J, et al. High Rate Triggers Increased Atrial Release of BMP10, A Biomarker for Atrial Fibrillation and Stroke, and BMP10 Affects Ventricular Cardiomyocytes. Circ Arrhythm Electrophysiol. 2025;18:e013834. doi: 10.1161/CIRCEP.125.013834

14. Lemme M, Ulmer BM, Lemoine MD, Zech ATL, Flenner F, Ravens U, Reichenspurner H, Rol-Garcia M, Smith G, Hansen A, et al. Atrial-like Engineered Heart Tissue: An In Vitro Model of the Human Atrium. Stem Cell Reports. 2018;11:1378–1390. doi: 10.1016/j.stemcr.2018.10.008

15. Krause J, Lemme M, Mannhardt I, Eder A, Ulmer B, Eschenhagen T, Stenzig J. Human-Engineered Atrial Tissue for Studying Atrial Fibrillation. Methods Mol Biol. 2022;2485:159–173. doi: 10.1007/978-1-0716-2261-2_11

16. Breckwoldt K, Letuffe-Breniere D, Mannhardt I, Schulze T, Ulmer B, Werner T, Benzin A, Klampe B, Reinsch MC, Laufer S, et al. Differentiation of cardiomyocytes and generation of human engineered heart tissue. Nat Protoc. 2017;12:1177–1197. doi: 10.1038/nprot.2017.033

17. Muller KF, Pan B, Ridder J, Schrapers J, Hansen J, Meier TM, Schulz C, Studemann T, Lemme M, Krause J, et al. Optogenetic tachypacing facilitates burst-induced arrhythmia in stem cell-derived human atrial engineered heart tissue. Europace. 2026;28. doi: 10.1093/europace/euag158

18. Morrell NW, Aldred MA, Chung WK, Elliott CG, Nichols WC, Soubrier F, Trembath RC, Loyd JE. Genetics and genomics of pulmonary arterial hypertension. Eur Respir J. 2019;53. doi: 10.1183/13993003.01899-2018

19. Packer M, Butler J, Ferreira JP, Siddiqi TJ, Januzzi JL, Jr., Sattar N, Maldonado SG, Panova-Noeva M, Prochaska JH, Sumin M, et al. Coordinated expression of BMP10/ALK1/endoglin-proteins that drive embryonic cardiac and vascular morphogenesis-in patients with heart failure: The EMPEROR Program. Eur J Heart Fail. 2025;27:1737–1751. doi: 10.1002/ejhf.3764

20. Steimle JD, Grisanti Canozo FJ, Park M, Kadow ZA, Samee MAH, Martin JF. Decoding the PITX2-controlled genetic network in atrial fibrillation. JCI Insight. 2022;7. doi: 10.1172/jci.insight.158895

21. 21. van den Berg NWE, Kawasaki M, Berger WR, Neefs J, Meulendijks E, Tijsen AJ, de Groot JR. MicroRNAs in Atrial Fibrillation: from Expression Signatures to Functional Implications. Cardiovasc Drugs Ther. 2017;31:345–365. doi: 10.1007/s10557-017-6736-z

22. Thomas AM, Cabrera CP, Finlay M, Lall K, Nobles M, Schilling RJ, Wood K, Mein CA, Barnes MR, Munroe PB, et al. Differentially expressed genes for atrial fibrillation identified by RNA sequencing from paired human left and right atrial appendages. Physiol Genomics. 2019;51:323–332. doi: 10.1152/physiolgenomics.00012.2019

23. Watanabe H, Darbar D, Kaiser DW, Jiramongkolchai K, Chopra S, Donahue BS, Kannankeril PJ, Roden DM. Mutations in sodium channel beta1- and beta2-subunits associated with atrial fibrillation. Circ Arrhythm Electrophysiol. 2009;2:268–275. doi: 10.1161/CIRCEP.108.779181

24. Yamada N, Asano Y, Fujita M, Yamazaki S, Inanobe A, Matsuura N, Kobayashi H, Ohno S, Ebana Y, Tsukamoto O, et al. Mutant KCNJ3 and KCNJ5 Potassium Channels as Novel Molecular Targets in Bradyarrhythmias and Atrial Fibrillation. Circulation. 2019;139:2157–2169. doi: 10.1161/CIRCULATIONAHA.118.036761

25. Burnham HV, Cizauskas HE, Barefield DY. Fine tuning contractility: atrial sarcomere function in health and disease. Am J Physiol Heart Circ Physiol. 2024;326:H568–H583. doi: 10.1152/ajpheart.00252.2023

26. Doehner W, Boriani G, Potpara T, Blomstrom-Lundqvist C, Passman R, Sposato LA, Dobrev D, Freedman B, Van Gelder IC, Glotzer TV, et al. Atrial fibrillation burden in clinical practice, research, and technology development: a clinical consensus statement of the European Society of Cardiology Council on Stroke and the European Heart Rhythm Association. Europace. 2025;27. doi: 10.1093/europace/euaf019

27. Pundi K, Gandotra C, Sanders W, Senatore F, Andrade JG, Gibson CM, Kirchhof P, Lubitz SA, Piccini J, Russo A, et al. Considerations for using atrial fibrillation burden as a surrogate endpoint: A report from the Cardiovascular Sciences Research Consortium. Am Heart J. 2026;301:107516. doi: 10.1016/j.ahj.2026.107516

28. Seibertz F, Rubio T, Springer R, Popp F, Ritter M, Liutkute A, Bartelt L, Stelzer L, Haghighi F, Pietras J, et al. Atrial fibrillation-associated electrical remodelling in human induced pluripotent stem cell-derived atrial cardiomyocytes: a novel pathway for antiarrhythmic therapy development. Cardiovasc Res. 2023;119:2623–2637. doi: 10.1093/cvr/cvad143

29. B. P. Atrial Fibrillation Modelling and Targeted DNA Methylation Editing in Human Engineered Heart Tissue-Based Disease Models. In: Electronic Dissertations of the State and University Library Hamburg (ediss): University of Hamburg; 2022.

