## Supplemental Material for "Dynamic BMP10 Release Reflects Atrial Fibrillation Burden in Human Atrial Engineered Heart Tissue"

### Supplemental Methods

#### RNA Isolation and Quantification

Snap-frozen aEHTs were homogenized in TRIzol reagent (Thermo Fisher Scientific) and processed using a phenol-chloroform extraction protocol. Chloroform was added to each homogenized sample, followed by vortexing and centrifugation at 12,000 g and 4°C for 15 minutes. The aqueous phase was transferred to a new tube, mixed with 70% ethanol, and loaded onto RNeasy Mini spin columns (Qiagen). RNA purification was performed according to the manufacturer's protocol, including on-column DNase digestion using the RNase-Free DNase Set (Qiagen). RNA was eluted in 15 µL RNase-free water and stored at −80°C until downstream use.

RNA concentration and purity were assessed using a µDrop Duo Plate and a Multiskan SkyHigh Microplate Spectrophotometer. For each sample, 1 µL RNA was measured in duplicate. RNase-free water was used as blank control. Absorbance was measured at 230, 260, and 280 nm, and RNA concentration as well as A260/A280 and A260/A230 ratios were calculated using the instrument software.

#### cDNA Synthesis

cDNA was synthesized from 200 ng total RNA using the High Capacity cDNA Reverse Transcription Kit (Applied Biosystems/Thermo Fisher Scientific). RNA was diluted with RNase-free water to a final sample volume of 10 µL. For each reaction, 10 µL cDNA master mix was added to 10 µL RNA sample, resulting in a final reaction volume of 20 µL.

The cDNA master mix per reaction contained 2 µL 10× RT buffer, 0.8 µL 25× dNTP mix, 2 µL 10× RT random primers, 1 µL MultiScribe Reverse Transcriptase, 1 µL RNase inhibitor, and 3.2 µL nuclease-free water. Reverse transcription was performed in a MiniAmp Thermal Cycler using the following program: 10 minutes at 25°C, 120 minutes at 37°C, and 5 minutes at 85°C. Synthesized cDNA was stored at −20°C.

#### Quantitative PCR Reaction Setup and Cycling Conditions

All cDNA used for qPCR was diluted 1:12 in nuclease-free water. qPCR reactions were performed in a final reaction volume of 10 µL, consisting of 4 µL diluted cDNA and 6 µL primer/SYBR Green master mix. The primer/SYBR Green master mix contained 5 µL

PowerTrack SYBR Green Master Mix (Applied Biosystems/Thermo Fisher Scientific), 0.5 µL primer mix, and 0.5 µL nuclease-free water per reaction.

Reactions were pipetted onto MicroAmp Optical 96-Well Reaction Plates and sealed with MicroAmp Optical Adhesive Film. qPCR was performed on a QuantStudio 5 Real-Time PCR Instrument. Samples were measured in technical triplicates. No-template controls were included for each primer pair, and water controls from cDNA synthesis were included.

The qPCR cycling protocol consisted of 2 minutes at 50°C and 10 minutes at 95°C, followed by 40 cycles of 15 seconds at 95°C and 1 minute at 60°C. Melt curve analysis was performed after amplification using 15 seconds at 95°C, 1 minute at 60°C, and 1 second at 95°C. Ct values were exported from QuantStudio Design and Analysis Software.

**qPCR Primer Design and In-Silico Validation**

qPCR primers were designed or selected using NCBI Primer-BLAST. Candidate primer pairs were screened against human RefSeq mRNA sequences and the Homo sapiens representative genome database. Primer properties were assessed using IDT OligoAnalyzer, including melting temperature, GC content, hairpin formation, self-dimerization, and hetero-dimerization. Primer sequences, amplicon lengths and validation details are listed in Supplementary Table 1.

For *CACNA1G*, genomic in-silico PCR predicted a 253 bp product at the *CACNA1G* genomic locus. For *KCNJ5* and *POLR2A*, only larger genomic products were predicted and considered low risk under qPCR conditions. For *PUM1*, no additional genomic target was detected. RNA samples were treated with DNase before cDNA synthesis to reduce genomic DNA contamination.

**BMP10 ELISA**

BMP10 concentrations in conditioned medium were quantified as described previously using the Human BMP-10 ELISA Kit PicoKine EK1808 (BosterBio, Pleasanton, CA, USA) according to the manufacturer’s instructions. Samples were diluted 1:10-1:50. Absorbance was measured using a Multiskan SkyHigh Microplate Spectrophotometer. BMP10 concentrations were calculated from the standard curve and are reported as ng/mL.

As the conditioned medium was collected every 2 or 3 days, the ELISA results were normalized to a 24 h release to allow for time-point comparison. Additionally, the medium used for feeding the aEHTs was confirmed to contain no measurable BMP10 using the same assay.

**Bulk RNA Sequencing and Bioinformatic Analysis**

Total RNA from control, intermittent 4 h/2 d (~10% burden), and continuous 24 h/day (100% burden) optogenetically paced aEHTs was used for bulk RNA sequencing. Libraries were prepared using the mRNA Library Prep Kit (Watchmaker Genomics).

Raw sequencing reads were processed with fastp (v0.23.2) using default parameters to remove sequencing adapter-derived sequences and low-quality reads. Mismatched base pairs in overlapping paired-end read regions were corrected using the fastp --correction option. Processed reads were aligned to the human reference assembly GRCh38.110 using STAR (v2.7.10a). Gene-level counts were generated using the STAR GeneCounts option (v2.7.10a).

To account for batch effects, count data were corrected using ComBat-Seq from the sva package (v3.54.0). Differential expression analysis was performed using DESeq2 (v1.42.0) based on the batch-corrected count matrix. Pairwise comparisons were performed between unpaced control, ~10% burden and 100% burden groups. Differentially expressed genes were defined by an absolute log2-transformed fold change ≥1 and a false discovery rate ≤0.1.

Overrepresentation analysis and gene set enrichment analysis were performed using clusterProfiler (v4.10.0). Gene sets included Gene Ontology terms and selected Molecular Signatures Database subsets, including Biocarta, KEGG, Reactome, WikiPathways, and Hallmark gene sets. Gene Ontology Biological Process gene sets were used for the normalized enrichment score heatmap.

For directional remodeling analyses, cumulative absolute log2 fold-change scores were calculated for genes across the control-to-intermediate and intermediate-to-continuous comparisons. Genes with the highest cumulative log2 fold-change scores and nominal *p* values ≤0.05 in both comparisons were selected for visualization. Volcano plots were generated from pairwise DESeq2 differential expression results using log2 fold change and adjusted *p* values. Heatmaps were generated from log1p transformed batch-corrected counts for selected gene sets.

### Supplementary Figures and Tables

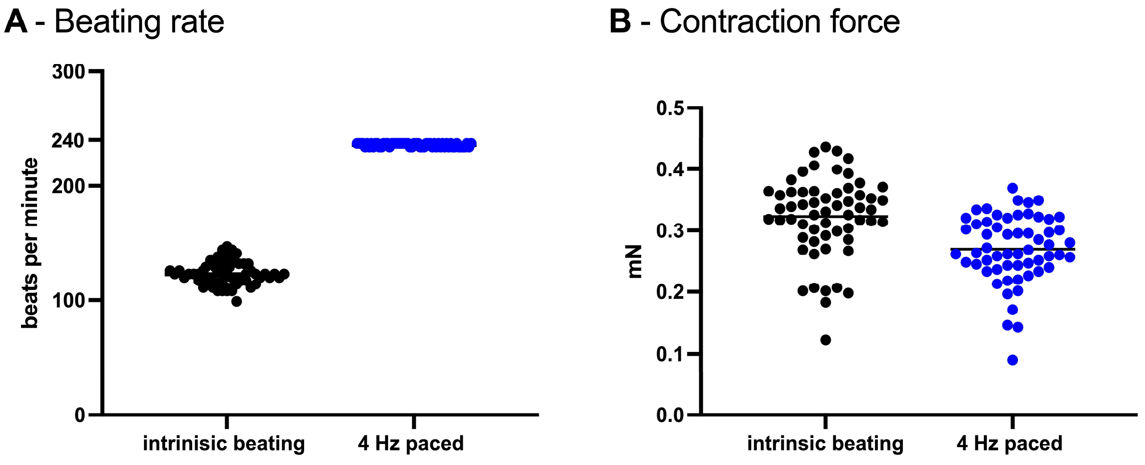


**Supplementary Figure 1 – Beating rate and contraction force during pacing**
**(A)** Beating rate of transduced aEHTs under intrinsic conditions and same aEHT during optical pacing at 4 Hz (470 nm light stimulus). All tissues followed the pacing stimulus, exhibiting a frequency shift from intrinsic rates (~2 Hz) to the applied high-rate pacing frequency of 4 Hz (n = 60). **(B)** Contractile force of aEHTs under intrinsic conditions and during optical pacing (n = 60). Each data point represents one aEHT; horizontal lines indicate mean values.

**A**

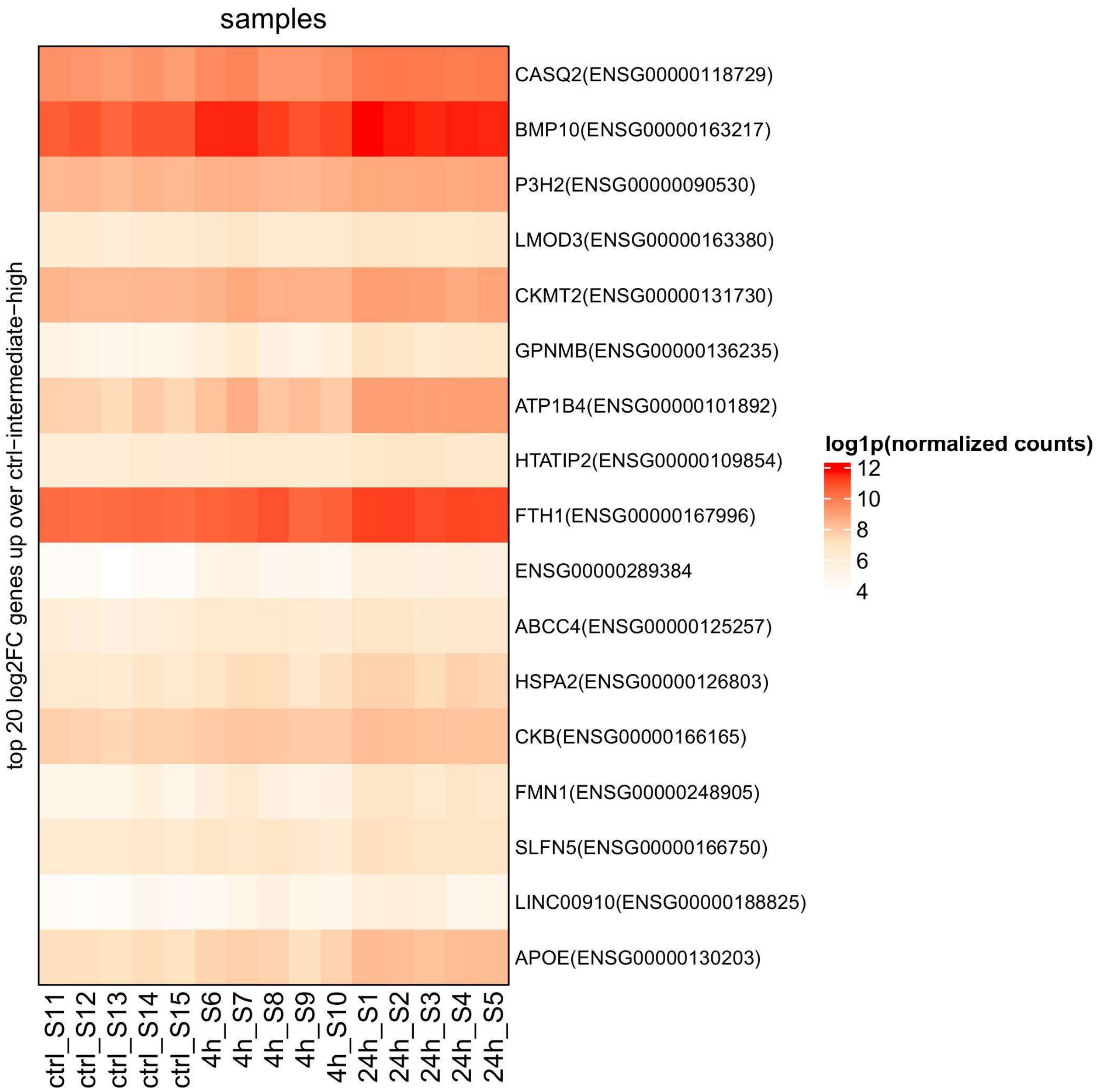
samples
top 20 log2FC genes up over ctrl−intermediate−high
CASQ2(ENSG00000118729)
BMP10(ENSG00000163217)
P3H2(ENSG00000090530)
LMOD3(ENSG00000163380)
CKMT2(ENSG00000131730)
GPNMB(ENSG00000136235)
ATP1B4(ENSG00000101892)
HTATIP2(ENSG00000109854)
FTH1(ENSG00000167996)
ENSG00000289384
ABCC4(ENSG00000125257)
HSPA2(ENSG00000126803)
CKB(ENSG00000166165)
FMN1(ENSG00000248905)
SLFN5(ENSG00000166750)
LINC00910(ENSG00000188825)
APOE(ENSG00000130203)
ctrl_S11
ctrl_S12
ctrl_S13
ctrl_S14
ctrl_S15
4h_S6
4h_S7
4h_S8
4h_S9
4h_S10
24h_S1
24h_S2
24h_S3
24h_S4
24h_S5
log1p(normalized counts)
12
10
8
6
4

**B**

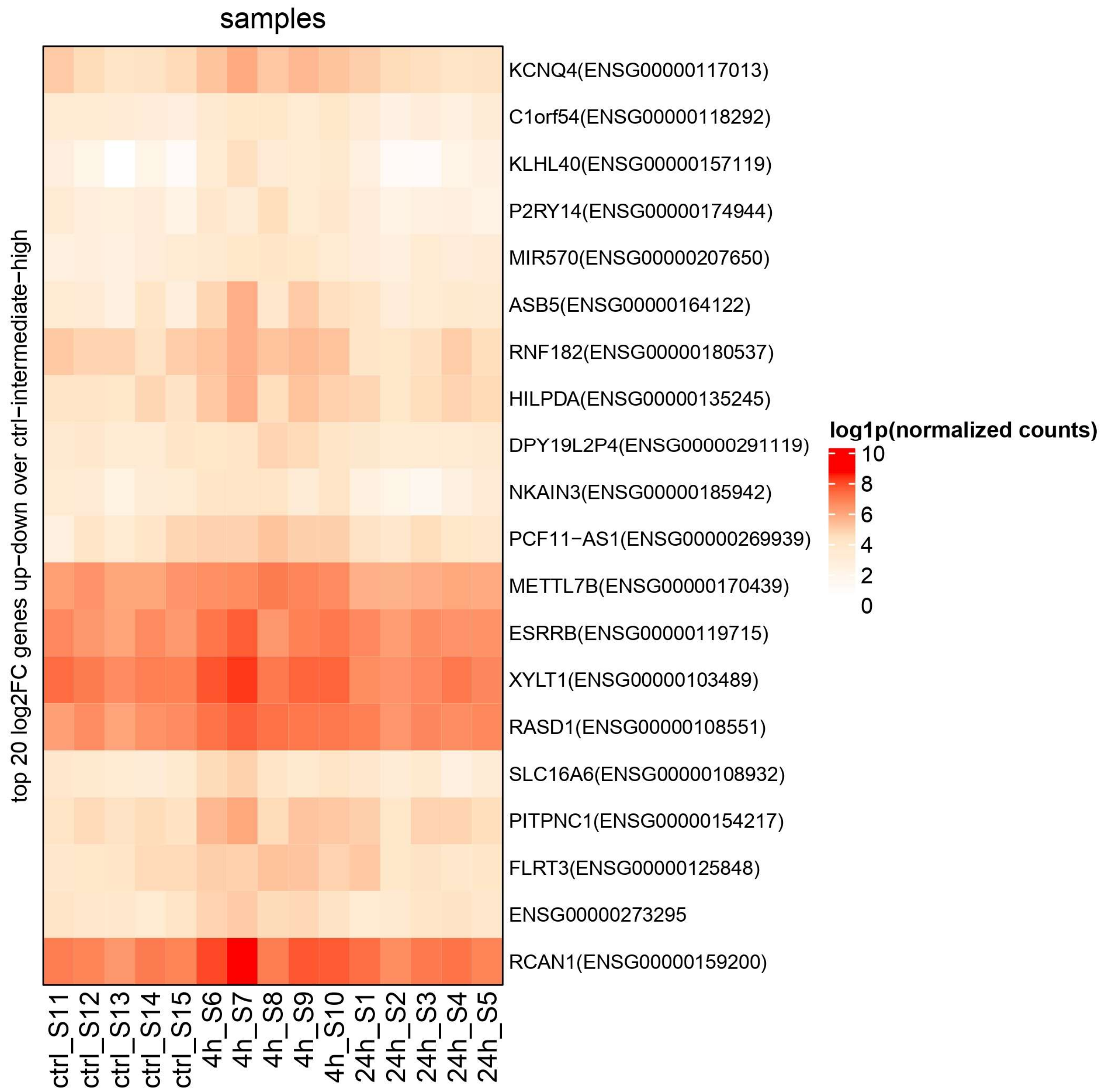


**Supplementary Figure 2 - Directional heatmaps of burden-associated transcriptional patterns**
Heatmaps showing selected transcripts with directional expression patterns across unpaced control, ~10% burden and 100% burden. (A) Top transcripts with monotonic upregulation across control-intermediate-high burden conditions, highlighting *BMP10* as part of the burden-associated upregulated transcriptional response. (B) Top transcripts with an up-down pattern across control-intermediate-high burden conditions, illustrating non-linear burden-dependent transcriptional regulation. Heatmaps show log1p-transformed normalized counts from batch-corrected RNA sequencing data. Genes were selected based on highest cumulative absolute log2 fold-change across the control-to-intermediate and intermediate-to-continuous comparisons with nominal *p* values ≤0.05 in both comparisons.

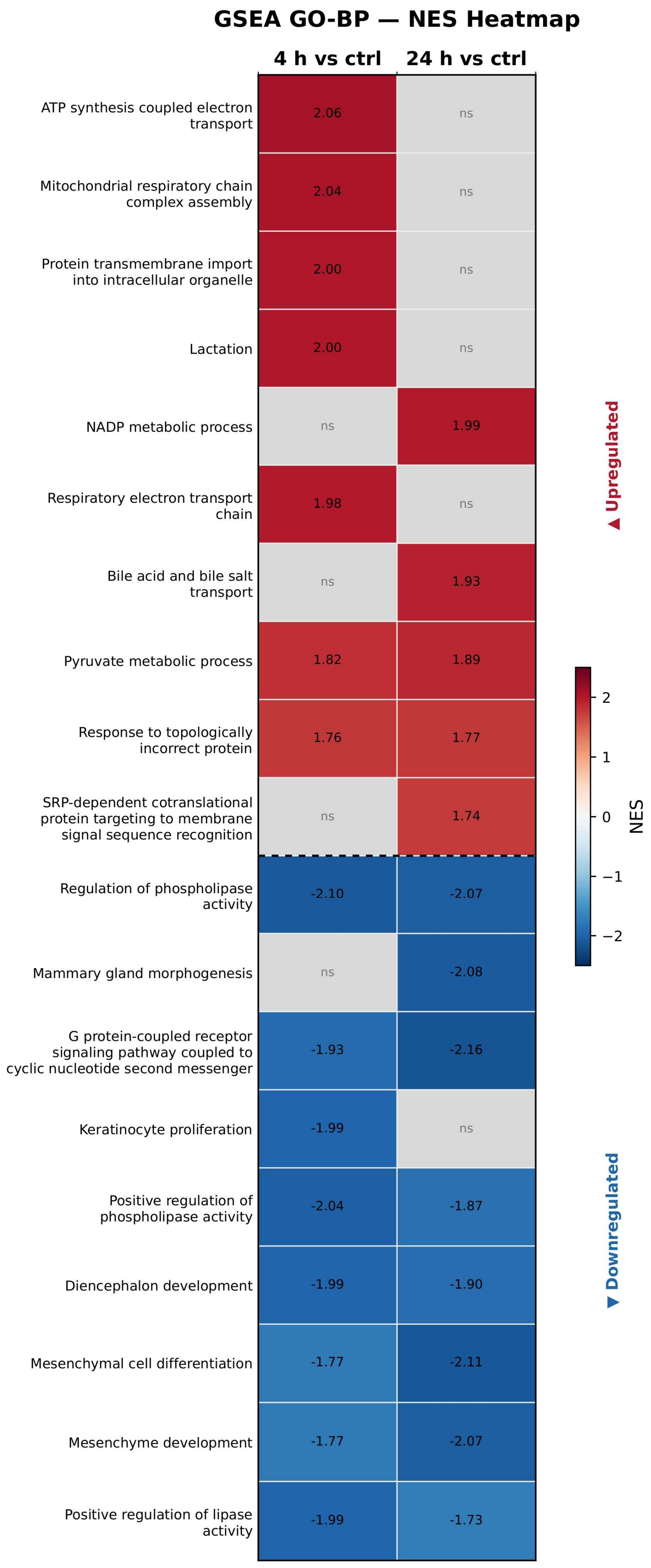
GSEA GO-BP — NES Heatmap
4 h vs ctrl
24 h vs ctrl
ATP synthesis coupled electron transport
2.06
ns
Mitochondrial respiratory chain complex assembly
2.04
ns
Protein transmembrane import into intracellular organelle
2.00
ns
Lactation
2.00
ns
NADP metabolic process
ns
1.99
Respiratory electron transport chain
1.98
ns
Bile acid and bile salt transport
ns
1.93
Pyruvate metabolic process
1.82
1.89
Response to topologically incorrect protein
1.76
1.77
SRP-dependent cotranslational protein targeting to membrane signal sequence recognition
ns
1.74
Regulation of phospholipase activity
-2.10
-2.07
Mammary gland morphogenesis
ns
-2.08
G protein-coupled receptor signaling pathway coupled to cyclic nucleotide second messenger
-1.93
-2.16
Keratinocyte proliferation
-1.99
ns
Positive regulation of phospholipase activity
-2.04
-1.87
Diencephalon development
-1.99
-1.90
Mesenchymal cell differentiation
-1.77
-2.11
Mesenchyme development
-1.77
-2.07
Positive regulation of lipase activity
-1.99
-1.73
▲ Upregulated
▼ Downregulated
NES
2
1
0
−1
−2

**Supplementary Figure 3 - ~10% and 100% burden induce distinct transcriptional changes in aEHTs**

Gene set enrichment analysis of Gene Ontology Biological Process terms was performed on bulk RNA sequencing data from human aEHTs after ~10% (4 h/2 d) or 100% burden (24 h/day) for 18 days, each compared with unpaced controls. The heatmap shows normalized enrichment scores (NES) for the top positively and negatively enriched terms.

**Supplementary Table 1 - Primers used for qPCR**

*POLR2A* and *PUM1* served as reference genes.

| Gene | Forward primer (5'-3') | Reverse primer (5'-3') | RefSeq / target transcript |
|---|---|---|---|
| *BMP10* | ACGCAGGATTCAGCCAAGGTG | ACCTCTTCATGGTGAGGAATGGAC | NM_014482.3 |
| *PITX2* | GTGTGGACCAACCTTACGGAAG | CGAAGCCATTCTTGCATAGCTCG | NM_000325.6 |
| *ENG* | CGGTGGTCAATATCCTGTCGAG | AGGAAGTGTGGGCTGAGGTAGA | NM_001114753.3 |
| *CASQ2* | GACGACTTTCCTCTGCTCGT | CCCCAATCTGTGGCCTGAAT | NM_001232.4 |
| *MYH7* | ACCAACCTGTCCAAGTTCCG | CCTCATTCAAGCCCTTCGTG | NM_000257.4 |
| *MYH6* | GCTCACCTACCAGACAGAGG | GCAAGAGTGAGGTTCCCGAG | NM_002471.4 |
| *CACNA1G* | TTCACCGCAGTCTTTCTGGCTG | TGACGGAGATGAGCACCAACAG | NM_018896.5 |
| *SCN1B* | CTGAGACCTTCACCGAGTGG | TCATAGCGCAGGATCTTGACA | NM_001037.5 |
| *KCNJ5* | GTGAAGGGACAGGATGGCTTA | GCTGGGATGTTGTTGAGATCG | NM_000890.5 |
| *POLR2A* | CTGCCAACATGACCTTTGCG | TGTGCCGTTCCACCTTATAGC | NM_000937.5 |
| *PUM1* | TTGAGTTTATTCCTTCAGACCAGC | GTGTGGATAAGGCAAATACCTGTC | NM_001020658.2 / NM_014676.3 |

This document was created with the Win2PDF “Print to PDF” printer available at

https://www.win2pdf.com

This version of Win2PDF 11 is for evaluation and non-commercial use only.
Visit https://www.win2pdf.com/trial/ for a 30 day trial license.

This page will not be added after purchasing Win2PDF.

Win2PDF Overview: https://www.win2pdf.com/video/overview/

Purchase Win2PDF: https://www.win2pdf.com/purchase/
